# Ecological selection for small microbial genomes along a temperate-to-thermal soil gradient

**DOI:** 10.1101/276584

**Authors:** Jackson W. Sorensen, Taylor K. Dunivin, Tammy C. Tobin, Ashley Shade

**Affiliations:** Department of Microbiology and Molecular Genetics, Michigan State University, East Lansing MI 48840 USA; Environmental and Integrative Toxicological Sciences, Michigan State University, East Lansing MI 48840; Department of Biology, Susquehanna University, Selinsgrove, PA 17870 USA; Department of Plant, Soil and Microbial Sciences; Program in Ecology, Evolutionary Biology and Behavior; and the Plant Resilience Institute, Michigan State University, East Lansing, MI 48840

**Keywords:** microbial ecology, genome size, genome reduction, thermophile, Centralia, coal seam fire, metagenome, disturbance, extreme environment

## Abstract

Small bacterial and archaeal genomes provide insights into the minimal requirements for life^1^ and seem to be widespread on the microbial phylogenetic tree^2^. We know that evolutionary processes, mainly selection and drift, can result in microbial genome reduction ^3,4^. However, we do not know the precise environmental pressures that constrain genome size in free-living microorganisms. A study including isolates ^5^ has shown that bacteria with high optimum growth temperatures, including thermophiles, often have small genomes ^6^. It is unclear how well this relationship may extend generally to microorganisms in nature ^7,8^, and in particular to those microbes inhabiting complex and highly variable environments like soil ^3,6,9^. To understand the genomic traits of thermally-adapted microorganisms, here we investigated bacterial and archaeal metagenomes from a 45°C gradient of temperate-to-thermal soils overlying the ongoing Centralia, Pennsylvania (USA) coal seam fire. There was a strong relationship between average genome size and temperature: hot soils had small genomes relative to ambient soils (Pearson’s r = −0.910, p < 0.001). There was also an inverse relationship between soil temperature and cell size (Pearson’s r = −0.65, p = 0.021), providing evidence that cell and genome size in the wild are together constrained by temperature. Notably, hot soils had different community structures than ambient soils, implicating ecological selection for thermo-tolerant cells that had small genomes, rather than contemporary genome streamlining within the local populations. Hot soils notably lacked genes for described two-component regulatory systems and antimicrobial production and resistance. Our work provides field evidence for the inverse relationship between microbial genome size and temperature requirements in a diverse, free-living community over a wide range of temperatures that support microbial life. Our findings demonstrate that ecological selection for thermophiles and thermo-tolerant microorganisms can result in smaller average genome sizes *in situ*, possibly because they have small genomes reminiscent of a more ancestral state.

## Main text

Genome streamlining is a reduction in genome size to increase cellular efficiency, and it evolves by means of selection^3^. A comparative analysis of changes in microbial genomes sizes with optimal growth temperature found a negative relationship that was independent of phylogeny and environment ^6^. This led to the conclusion that thermophiles are examples of free-living microorganisms subject to genome streamlining ^6,10,11^. These results were exciting because they suggested that high temperature can select on genome size, providing insights into environmental conditions that may propel efficiency. For the comparative analysis ^6^ and cited studies therein, temperature optimum, genome size, 16S rRNA gene sequences, and habitat were available for a curated collection 115 bacterial and archaeal isolates ^5,12^. Given biases of cultivation ^13^, an outstanding question was whether the relationship between growth temperature and genome size would prove to be general for wild microbial communities.

Fortuitously, the fire-impact gradient at the Centralia ecosystem provides an opportunity to investigate relationships between temperature and microbial genome traits. Centralia, Pennsylvania is the site of a slow-burning, near-surface coal seam fire that ignited in 1962. The heat from the fire vents through overlying soils, causing surface soil temperatures to reach as high as > 400°C ^14^, but more recently in the range of 40 - 75°C ^15,16^. However, the soils in Centralia were previously temperate, with no known exposure to prolonged high temperatures. Therefore, Centralia offers an interesting model for the examining the eco-evolution of microbial communities ^17^.

We recently used 16S rRNA gene amplicon sequencing to assess compositional changes in Centralia soil microbial communities along an ambient-to-thermal temperature gradient ^16^. Surface soils overlying the coal seam fire were collected to include soils that were hot from fire (“fire-affected”), soils that were previously hot but had since recovered to ambient temperatures (“recovered”) and reference soils that had never been impacted by the fire. As expected, fire-affected soils had starkly different community structure from ambient soils. However, after the fire advanced, soils reasonably recovered towards reference community structure. This suggested a considerable capacity of soil microbiomes for resilience, even after exposure to a severe and unanticipated stressor, and prompted us to next ask what microbial attributes underlay the observed changes in community structure in fire-affected soils.

Moving forward, we assessed average genome size along the Centralia fire gradient (**Table S1**). From twelve metagenomes (six fire-affected, five recovered, and one reference), we used MicrobeCensus ^18^ to calculate average genome size across a soil temperature range of 45 °C. Average genome sizes were negatively and strongly correlated with temperature (**Figure 1A**, Pearson’s r = −0.910, p < 0.001). In addition to MicrobeCensus, we used three other distinct and complementary methods to assess changes in genome size with soil temperature and found them all to be in agreement **(Figure S1).** To the best of our knowledge, this is the first report of decreases in genome size across an *in situ* temperature gradient that supports the broad range of physiological requirements from mesophiles to thermophiles.

**Figure 1.**
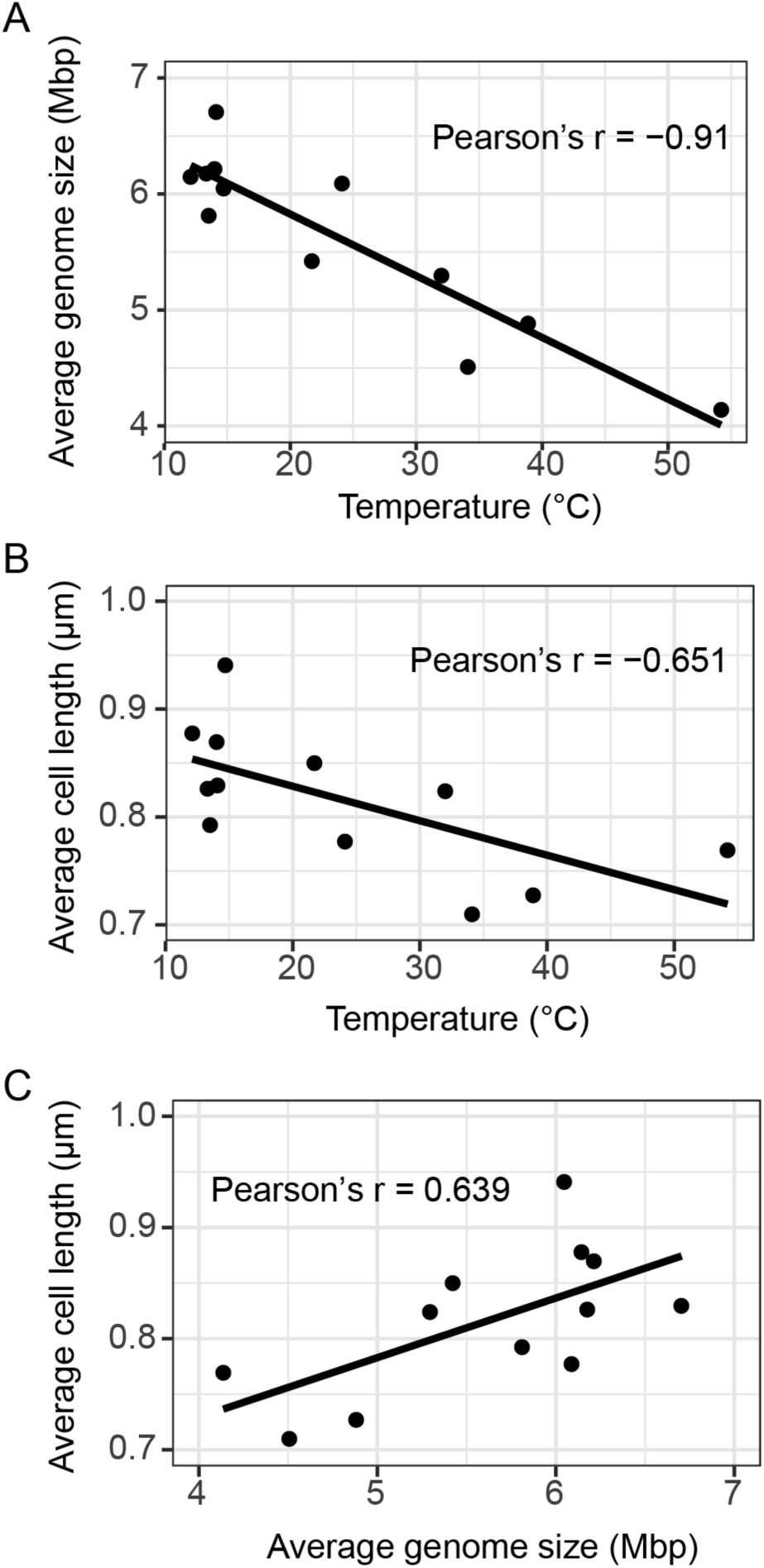
Changes in average genome and cell sizes across the soil temperature gradient in Centralia. (A) Average genome size in each metagenome was calculated using MicrobeCensus and plotted against site temperature. (B) Average cell length was measured from 44-910 cells from 3-9 replicate fields for each soil and plotted against soil temperature. (C) Average genome size had a direct relationship with average cell size.

We next compared the average genome sizes estimated from Centralia metagenomes to those from 22 publicly available soil metagenomes (**Figure 2, Table S2**). Generally, hot soils in Centralia had small genomes relative to other soils, while ambient soils in Centralia were closer to the average size observed among this set. Intriguingly, permafrost soils also harbored small average genomes and were comparable to the hottest Centralia sites. These results support comparably small genome sizes in Centralia soils and also provide a range of expected soil genome sizes more generally.

**Figure 2.**
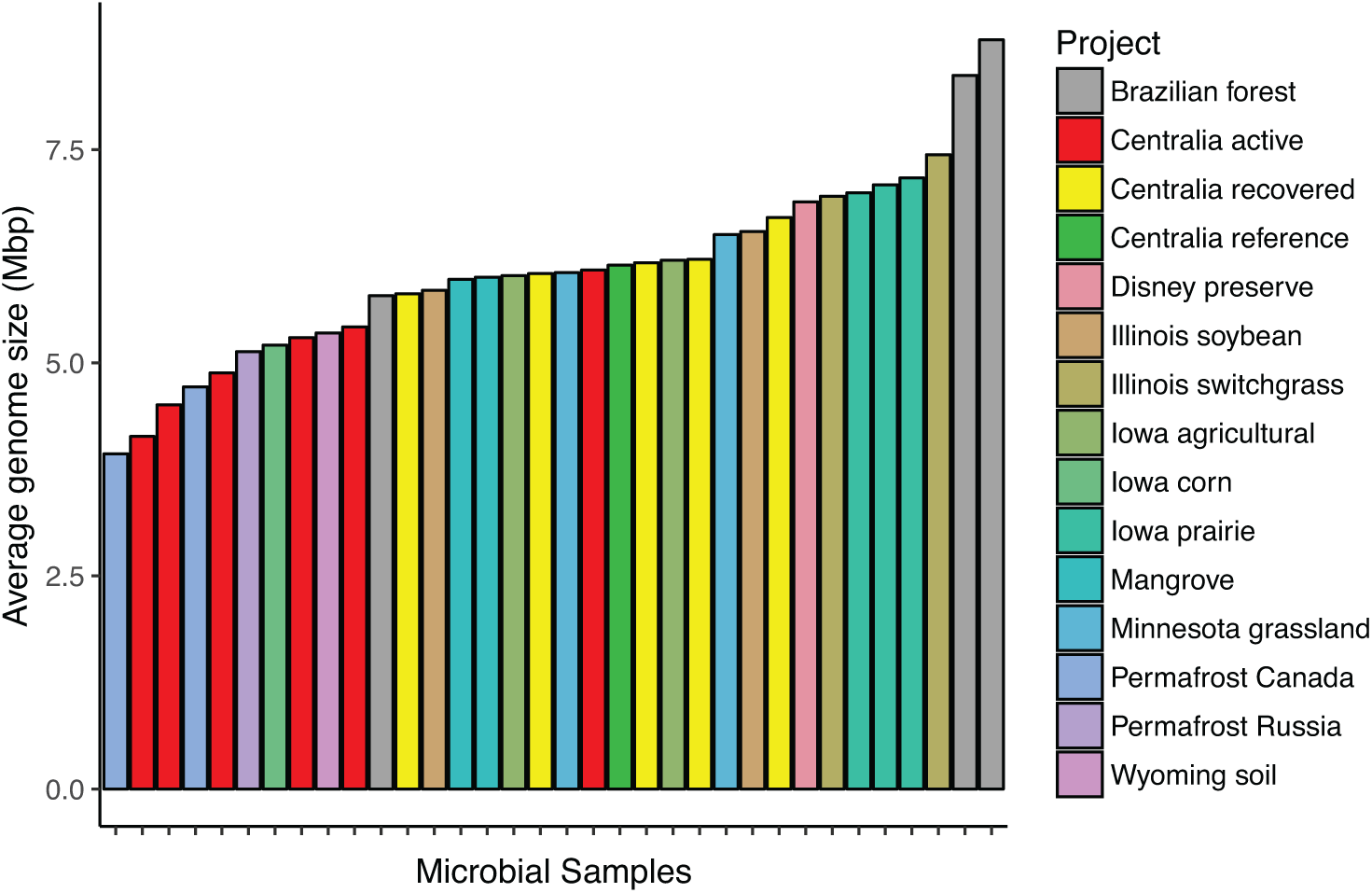
Average genome size in soil metagenomes, estimated using MicrobeCensus.^18^ Samples are ordered by average genome size and colored by sample location.

It was hypothesized that small cells may be selected to attain minimal cellular maintenance costs at high temperatures, and that small cells indirectly select for small genomes ^6^. Because we had microscope images from soil cell counts in Centralia^16^, we re-analyzed the images to extract size information. We found that average cell sizes were also negatively correlated with temperature (**Figure 1B**, Pearson’s r = −0.65, p =0.021). Accordingly, cell size had a direct relationship with genome size (**Figure 1C**, Pearson’s r = 0.64, p = 0.025). These results agree with reported *in situ* relationships between cell size and temperature in aquatic systems. For example, an experiment investigating a 6°C increase in water temperature confirmed that smaller cells with lower nucleotide content were selected at warmer temperatures ^7^, providing support that even slight warming may enrich for microorganisms with small genomes. An observational study of marine microbial genome size along a latitudinal gradient (10.7°C range) also supports this hypothesis ^8^. Our results extend the cell size-temperature trend to soils and also to a temperature range encompassing 45 °C.

To understand the selective outcomes of high temperature on the functions of these small genomes, we next asked if there were functional genes that were characteristically enriched or depleted with increasing temperature. We used shotgun metagenome annotations from the KEGG module (KM) database ^19^. KMs are groups of KEGG Orthologs (KOs) that represent complexes, functional sets, metabolic pathways, or signatures. Eighty-one percent of KOs detected in Centralia metagenomes were detected in all soils, and many patterns with temperature were attributable to changes in normalized KO abundance rather than in KO detection. In total, 284 (out of 541 detected; 52.50%) were correlated with temperature (**Figure 3**, **Table S3**).

**Figure 3.**
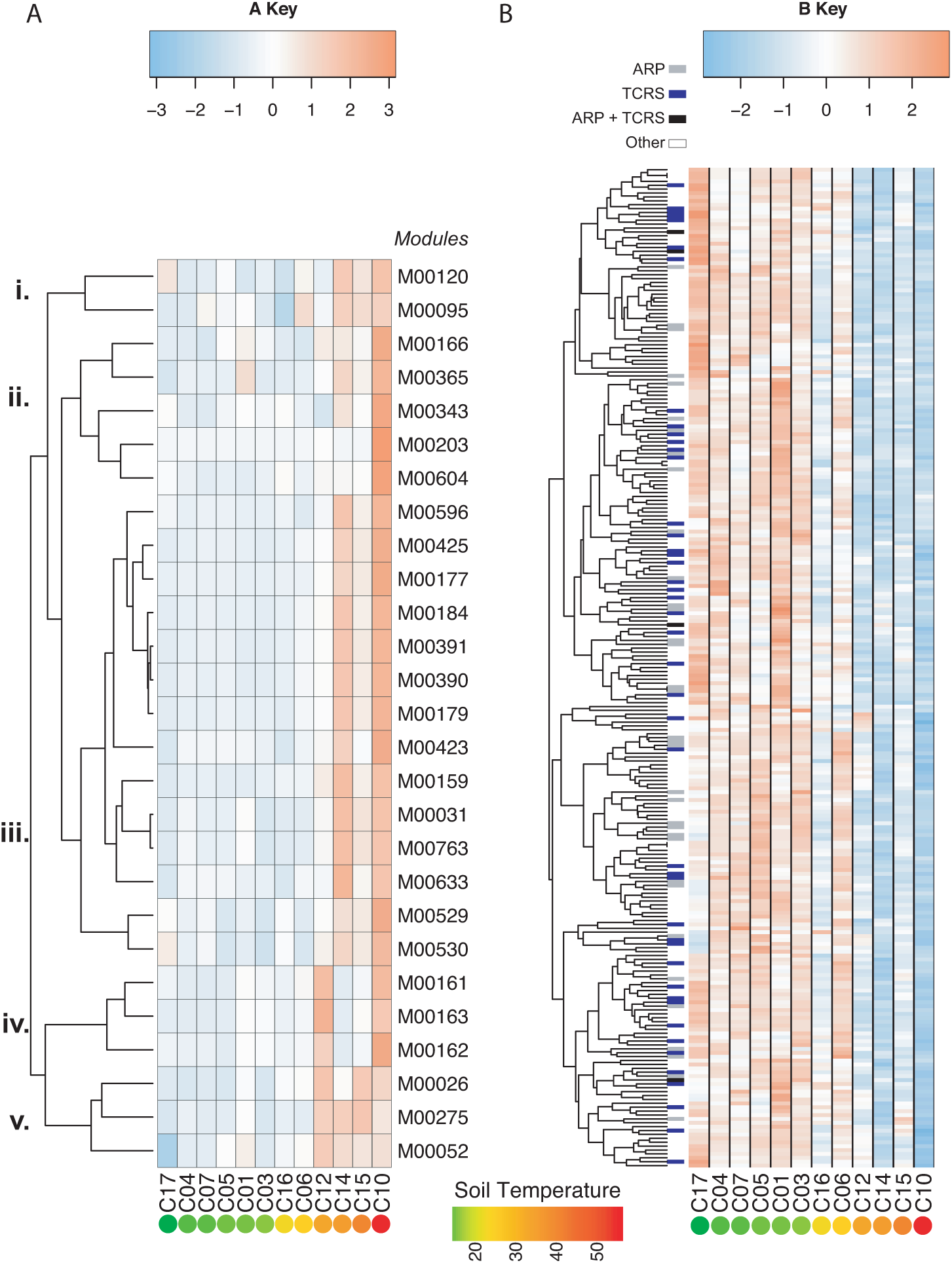
Heatmap of KEGG modules correlated with temperature (false discovery rate adjusted p-value < 0.05). Modules (rows) are centered and standardized across Centralia metagenomes (columns), with warm colors showing relative enrichment and cool colors showing relative depletion. Modules with significant relationships with temperature are shown. Sites are arranged by increasing temperature from left to right. (A) 27 KEGG modules were positively correlated with temperature (Pearson’s R range = 0.646 to 0.933). (B) 257 KEGG modules were negatively correlated with temperature (Pearson’s R range = −0.642 to −0.925). A third of the KEGG modules negatively correlated with temperature were either two-component regulatory systems (TCRS, blue dendrogram tips), antimicrobial resistance or production (ARP, gray tips), or both (black tips). Note differences in color gradient ranges across panels A and B.

Twenty-seven KMs were positively correlated with temperature (Pearson’s R > 0.656, false discovery rate adjusted p-value < 0.05; **Figure 3A**). Specifically, dissimilatory sulfate reduction (M00596), dissimilatory nitrate reduction (M00530) and denitrification (M00529) were enriched in hot soils (**Figure 3A**, *cluster iii*; **Figure 4A**). These are anaerobic processes aligned with known and expected environmental conditions in Centralia. Fire-affected soils from active vents have higher moisture than reference and recovered soils (Pearson’s r = 0.714, p < 0.01), which likely promote inundated and anaerobic microhabitats therein. Prior work in Centralia has indicated an importance of these metabolisms in hot soils, noting that sulfur, sulfate, nitrate and ammonium were commonly elevated at vents ^14,15^. These results also agree with observations of thermophile metabolisms in other terrestrial and geothermal environments, including a prevalence of denitrification and dissimilatory nitrate reduction ^20,21^, highly active nitrogen cycles in hot springs ^22^, and increased dissimilatory organic sulfur mineralization ^23^. Notably, these anaerobic KMs grouped in their response patterns with several archaeal proteins (**Figure 3A** *cluster iii*; Archaeal ribosome M00179, polymerase M00184, and exosome M00390). We also observed an increase in Crenarchaeota in fire-affected soils ^16^, an archaeal phylum that includes sulfate reducers ^24^. Additional results describing patterns and thresholds of KM enrichment with temperature are provided in Supporting Materials. Together, these data suggest that the pathways enriched in small genomes from hot soils offer functions attuned to the Centralia habitat.

**Figure 4.**
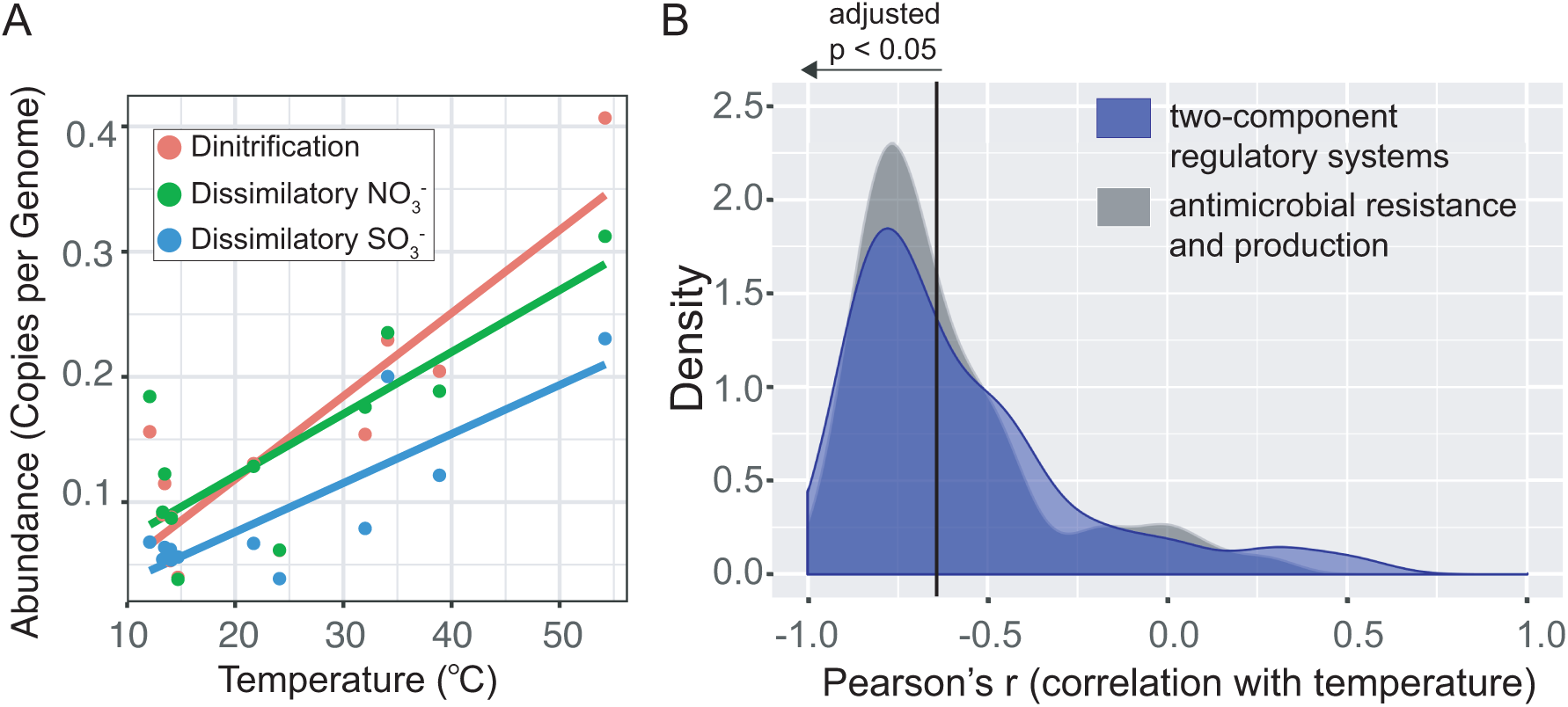
KEGG modules that had notable enrichments or depletions with temperature. (A) The median abundances of KEGG modules for denitrification (red), dissimilatory nitrate reduction (green) and dissimilatory sulfate reduction (blue) were all positively correlated with temperature. (B) Pearson’s correlation values for all detected modules classified as antibiotic resistance and production (gray density, n = 62 detected modules) or two-component regulatory systems (blue density, n = 89 detected modules). The black vertical line distinguishes correlation values that are significant at a false discovery rate adjusted p-value <0.05 (left), and all of these had a strong and negatively relationship with temperature. In total, there were 39 antimicrobial resistance and production modules and 49 two-component regulatory system modules that significantly decreased with temperature.

Temperature was negatively correlated with 257 KMs (47.5% out of 541 total KMs detected, Pearson’s R < −0.6, false discovery rate adjusted p-value < 0.05; **Figure 3B**). In general, these depleted KMs were detected across recovered soils and the reference soil. There were two noteworthy categories of KMs that were consistently depleted in hot soils: antimicrobial resistance and production and two component regulatory systems (**Figure 4B)**. Together, these two KM categories comprised 32.7% of KMs negatively correlated with temperature (84 out of 257). This trend was striking, but we also note that some KMs belonging to these categories had no relationships with temperature and that these KM categories were always detected in fire-affected soils.

Thirty-nine modules for antimicrobial production and resistance mechanisms were negatively correlated with temperature (**Figure 4B),** which agrees with a prior analysis of antibiotic resistance genes in this system ^25^. Among these modules were resistance to vancomycin, tetracycline, fluoroquinolone, aminoglycoside, nisin, erythromycin, streptomycin and beta-lactam, and several multidrug efflux pumps. The small genomes of host-associated symbionts often lack antimicrobial genes ^26^. However, the *Pelagibacter* clade, which is a model free-living population that has streamlined genomes, has a conserved multidrug transporter across sequenced genomes ^27^. It could be that thermophiles have fewer genes encoding resistance to described antimicrobials, as evidenced by the challenges inherent in developing specific selectable antibiotic resistance markers for thermophiles ^28,29^. A related consideration is that, like most databases, KEGG is biased towards genomes and annotations from fast-growing mesophiles and may have missed annotation of under-described thermophile antimicrobials. To clarify whether the observed decrease in antimicrobial production and resistance was due to unannotated novelty or a true deficit of these functions in thermal sites, annotation-independent methods could be used to identify antimicrobial-related biosynthetic gene clusters from Centralia metagenomes ^30,31^. In addition, functional screens of Centralia isolates could be performed for antibiotic production and resistances. If there is a true deficit in genes encoding antimicrobial production and resistance, it could be that the thermal conditions present a strong environmental filter that reduces competition among the populations tolerant of the heightened temperature. Our previous work reported decreased richness and phylogenetic diversity fire-affected Centralia soils ^16^, suggesting that there is a smaller pool of potential competitors inhabiting the hot soils.

Additionally, forty-nine detected two-component regulatory system modules were also negatively correlated with temperature (Pearson’s R < −0.6, **Figure 4B**). Two-component systems consist of a sensor kinase and a response regulator and allow for transcriptional responses to environmental stimuli ^32^. This simple regulatory system allows bacteria to respond to multiple stimuli: the involved genes duplicate, the sensors evolve sensitivity to additional stimuli, and additional genes are transcribed ^32,33^. Previous studies suggested that smaller genomes have fewer regulatory components ^34^, and this relationship is often observed in streamlined genomes ^3,8^. Our results agree with observations of generally less regulation with smaller genomes^4,11,27,35,36^ and also suggest that thermophiles may have lower regulatory needs. It has been proposed that thermophiles with “streamlined” genomes may be more likely to utilize global regulatory systems that mediate transcriptional responses to co-occurring environmental stimuli ^11^. The degree of environmental variability is also predicted to influence the relative benefit an organism gains from investing in sensing its environment ^37^. As a common case study in genome reduction, obligate endosymbionts are thought to have drifted towards small genomes in part because environmental conditions are stable and thus sensing requirements are minimal (e.g., ^3^). Furthermore, in Centralia, seasonal temperature fluctuations in fire-affected soils are equivalent to those in ambient soils (**Figure S2**), providing evidence that the soils experience similar environmental stability in temperature, albeit at different ranges. This suggests that small genomes are not necessarily conditional on very stable environments ^3^. Future work should investigate whether two-component regulatory systems are consistently less prevalent among thermophiles, and, if so, whether their absence is reminiscent of an ancestral state.

Our field study supports and reinforces cultivation-dependent observations that suggested bacteria and archaea with small genome sizes have higher growth temperatures ^6^. Because our study considers ecological section, as evidenced by the turnover in community membership between ambient and hot soils^16^, these data indicate that environmental microorganisms with relatively higher temperature requirements also are likely to have small genomes and cell sizes. Surprisingly, it also suggests that microbial populations inhabiting complex environments, like soils, may generally reflect similar overarching traits in genome size as those observed in laboratory studies, which are necessarily biased towards fast-growing organisms that often are of medical, industrial, or agricultural interest (e.g., ^38^). In addition, this work expands upon previous reports of smaller genomes with higher temperatures ^7,8^ to consider a range of *in situ* temperatures at which a variety of microbes compete in non-optimal conditions. For example, we would expect mesophiles growing near their upper temperature ranges and thermophiles growing near their lower temperature ranges to co-occur at some sites in Centralia. Therefore, these results are relevant to the experiences of many wild microorganisms that cope with dynamic environments.

Our results add evidence that supports both smaller genomes and cells, on average, with higher temperatures but also offer a key point of distinction. Though the taxa enriched in Centralia hot soils characteristically had smaller genomes and cells, there is no evidence for contemporary genome streamlining in Centralia. Rather, we suspect that these thermo-tolerant cells were resuscitated from the vast dormant pool in soil. This is supported by three lines of evidence. First, there was turnover in community membership across hot and ambient Centralia soils ^16^, providing evidence against contemporary streamlining within local lineages. Second, there was striking comparability in average genome size of hot Centralia soils to ancient permafrost soils, which largely contain an inactive and very old dormant pool. Third, many other studies have described thermophile persistence and resuscitation from non-thermal environments, suggesting that these lineages are widespread but typically inactive ^21,39,43^. Therefore, we posit that Centralia small genomes are characteristic of an ancestral trait of previously dormant thermophiles in the soil and not the outcome of genome streamlining.

In conclusion, we found a strong negative relationship between average microbial genome size and temperature in Centralia soils along a mesophile-to-thermophile gradient, spanning 45°C. We also found that cells were smaller in hot soils, supporting the hypothesis that thermo-tolerant bacteria have smaller cell size, which indirectly selects for small genomes^6^. By KEGG annotations, Centralia metagenomes at hot temperatures were best defined by what they lacked rather than enriched modules of distinctive metabolisms. Specifically, environmental sensing mechanisms, such as two-component regulatory systems, and antimicrobial production and resistance mechanisms were in lower abundance in hot soils. In addition, there were a few modules enriched at high temperatures that met expectations for the hot anaerobic environment at active vents, including nitrogen and sulfur metabolism. Our results show that the relationship that was observed between growth temperature and genome size for cultivable isolates also holds true in a complex, *in situ* microbial community that inhabits a complex and variable soil environment. We suggest that, for thermo-tolerant organisms, the relationship between temperature and genome size indicates the precursory microbial condition of small genomes, reminiscent of ancient lineages, rather than contemporary genome streamlining.

## Materials and Methods

### DNA extraction and metagenome sequencing

DNA for metagenome sequencing was manually extracted using a phenol chloroform extraction ^44^ and then purified using the MoBio DNEasy PowerSoil Kit (MoBio, Solana Beach, CA, USA) according the manufacturer’s instructions. Total DNA sequencing was performed on all 12 samples by the Department of Energy’s Joint Genome Institute (Community Science Project) using an Illumina HiSeq 2500. Libraries were prepared with a targeted insert size of 270 base pairs. Samples had between 19Gbp and 50Gbp of sequence data. Additional methodology details are provided in Supporting Materials.

### Quality control, assembly and annotation

Assembly was performed by the Joint Genome Institute according to their standard operating procedure (Supporting Materials). To use all sequencing data, we worked with assembled and unassembled reads processed by Integrated Microbial Genomes (IMG) using their standard annotation pipeline^45^. After comparing several annotation methods (Supporting Materials), we chose to use the KEGG Orthology database for analyzing the Centralia data due to its inherent structure and ability to integrate metabolic pathways. KEGG Ortholog (KO) abundances were relativized to the median abundance in each site of a set of 36 single copy genes published previously^46^ (see Supporting Materials). One single copy gene (K01519) was an outlier in 7 out of 12 samples as assessed by Grubb’s test for outliers and removed. We analyzed patterns in KEGG Modules (KMs)^19^, a set of manually defined functional units made up of multiple KOs. KM abundances were calculated based on the median abundance of their constituent KOs that were present in the metagenomes. KMs were included in analysis if 50% or more of their constituent KOs were identified in the dataset. Approximately one third of the open reading frames per sample were able to be annotated with KEGG (**Table S1**). As a caveat to the study, unannotated open reading frames can result from erroneous reads and mis-assemblies but also could be novel and or divergent genes critical for microbial processes. Thus, new annotations could impact the overarching patterns described here.

### Average genome and cell size

Average genome size was calculated from the quality filtered DNA sequences using MicrobeCensus (“run_microbe_census.y –n 2000000”), which estimates average genome size by calculating the percent of sampled reads that match to a set of single copy genes ^18^. We also used three additional methods to calculate average genome size (see Supporting Materials), and all were in agreement in revealing the negative relationship between temperature and average genome size. To calculate cell size, we re-analyzed microscope images previously used to count microbial cells for community size quantifications in the same soils ^16^. We hand-curated a debris-free subset from the images and measured 44 - 910 cells from 3 - 9 replicate fields for each soil. The major and minor axes of cells were measured using a FIJI macro in ImageJ (Version: 2.0.0-rc-65/1.51s Build: 961c5f1b7f). We found that cell size range and deviations (**Table S4**) were consistent with those previously reported ^48^.

### Comparisons with other soil metagenomes

All metagenomic data sets for comparison were obtained from MG-RAST ((http://metagenomics.anl.gov/). The MG-RAST database was searched with the following criteria: material = soil, sequence type = shotgun, public = true. The resulting list of metagenome data sets were ordered by number of base pairs (bp). Metagenomic data sets with the most bp were included if they were sequenced using Illumina (to standardize sequencing errors), had an available FASTQ file (for internal quality control), and contained < 30% low quality as determined by MG-RAST. Within high quality Illumina samples, priority for inclusion was given to projects with multiple samples. When a project had multiple samples, data sets with the greatest bp were selected. This search yielded 22 data sets from 12 locations and five countries (**Table S2**). Sequences from MG-RAST data sets were quality checked using FastQC (v0.11.3, ^49^ and quality controlled using the FASTX toolkit (fastq_quality_filter, “-Q33 −q 30 −p 50”). Average genome size for each dataset was calculated from the quality filtered DNA sequences using MicrobeCensus with default parameters.

### Statistical analyses

Statistics for the metagenome datasets were performed in the R environment for statistical computing^50^. The stats package was used for calculating Pearson’s correlations^50^. The outliers package ^51^ was used for identifying outlying KOs. The ggplot2 package was used for visualization^52^. Heat maps were created with heatmap2 from the gplots package^53^.

### Data and workflows

All analysis workflows are available on GitHub (ShadeLab/PAPER_SorensenInPrep). Metagenome data are available on IMG under the GOLD Study ID GS0114513.

## Acknowledgements

This research was supported by Michigan State University. Computational resources were provided by the Institute for Cyber-Enabled Research. Metagenome sequencing was supported by the Joint Genome Institute Community Science Project #1834. The work conducted by the U.S. Department of Energy Joint Genome Institute, a DOE Office of Science User Facility, is supported under Contract No. DE-AC02-05CH11231. We thank Keara L. Grady and Sang-Hoon Lee for technical support and Susanna Yeh (REU-ACRES NSF Grant # 1560168) for assistance with soil metagenome curation from MG-RAST and analyses.

## Contributions

AS and TCT conceived the study and conducted field work. JWS and TKD performed analyses, with direction and oversight by AS. JWS, AS and TKD contributed writing. All authors discussed results, and commented on and edited the manuscript.

## Supporting Figures

Figure S1. Complementary methods used to assess changes in average genome size across the soil temperature gradient in Centralia. (A) Odds ratios were calculated for 35 single-copy gene KEGG Orthologs in each site and plotted against site temperature. Reported correlation is between all single copy gene odds ratios and temperature, and all p < 0.001. (B) Average genome size in each site was calculated based on phylum level abundances from 16S rRNA gene amplicon data, using weighted average genome sizes of each phylum present in JGI IMG (accessed 19 June 2017, correlation p < 0.001). (C) Average MAG size at each site was calculated based on presence/absence of 104 MAGs (correlation p = 0.029).

Figure S2. Annual temperature fluctuations at three fire-affected (circles) and two ambient (triangles) Centralia sites, measured using *in situ* temperature loggers (HOBOs) that were buried 5 - 10 cm below the surface. Temperature loggers were deployed after the soils were collected for this study.

## Supporting Tables

Table S1. Sequence summary information for Centralia metagenomes. Soils were collected 03- 07 October 2014. Asterisks indicate that the site was actively venting at the time of soil collection.

Table S2. MG-RAST metadata for soil metagenomes used in this study.

Table S3. KEGG Modules significantly correlated with temperature (false-discovery-rate adjusted p-value <0.05)

Table S4. Cell size measurements from microscope images.

Table S5. Single-copy KEGG Orthologs’ odds ratios correlations with temperature.

Table S6. Lineage, completeness and contamination of Metagenome Assembled Genomes as estimated by CheckM

